# Development of a Tet-On inducible expression system for the anhydrobiotic cell line, Pv11

**DOI:** 10.1101/2020.05.29.123570

**Authors:** Shoko Tokumoto, Yugo Miyata, Kengo Usui, Ruslan Deviatiiarov, Takahiro Ohkawa, Sabina Kondratieva, Elena Shagimardanova, Oleg Gusev, Richard Cornette, Masayoshi Itoh, Yoshihide Hayashizaki, Takahiro Kikawada

## Abstract

The Pv11 cell line established from an African chironomid, *Polypedilum vanderplanki*, is the only cell line tolerant to complete desiccation. In Pv11 cells, a constitutive expression system for Pv11 cells was previously exploited and several reporter genes were successfully expressed. Here we report the identification of an effective minimal promoter for Pv11 cells and its application to the Tet-On inducible expression system. First, using a luciferase reporter assay, we showed that a 202 bp deletion fragment derived from the constitutively active 121-promoter functions in Pv11 cells as an appropriate minimal promoter with the Tet-On inducible expression system. The AcGFP1 protein was also successfully expressed in Pv11 cells using the inducible system. In addition to these reporter genes, avian myeloblastosis virus reverse transcriptase α subunit (AMV RTα), which is one of the most widely commercially available RNA-dependent DNA polymerases, was successfully expressed through the inducible expression system and its catalytic activity was verified. These results demonstrate the establishment of an inducible expression system in cells that can be preserved in the dry state and highlight a possible application to the production of large and complex proteins.

**Highlights:**

- A 202 bp-deletion fragment derived from a constitutively active promoter was identified as a minimal promoter in Pv11 cells.
- A Tet-On inducible expression system was developed for Pv11 cells using the minimal promoter.
- Typical reporter genes (GFP and luciferase) and an enzyme with complex structure, i.e. a viral reverse transcriptase, were successfully and inducibly expressed in Pv11 cells using the Tet-On system.

## 1. Introduction

Recombinant proteins have many applications in the biotechnology and biomedical industries. The production of recombinant proteins of sufficient quantity and quality for commercial applications relies on the selection of an appropriate combination of gene expression system and host cell [1]. Insect cells are one of the most common hosts for recombinant protein expression because they can produce larger and more complex proteins, whose biological activities are maintained, than is possible in bacterial cells [2].

Many gene expression systems for insect cells have been developed over recent years [3-5]. For example, insect cell-based expression systems are used for the commercial production of various veterinary and human vaccines [6]. Generally, insect cells are cultured at ambient temperature without a CO_2_ incubator, and thus are more cost-effective hosts than mammalian cells. To date, more than 1,100 insect cell lines have been established according to the ExPASy Cellosaurus database (http://web.expasy.org/cellosaurus). However, relatively few of these are regularly used as protein expression systems, and the potential of other cell lines for protein expression has not been fully explored, especially in the case of non-model insect cell lines. Thus, using a cell line with specific features (e.g., low-temperature tolerance [7]) would benefit the development of innovative biotechnologies for protein production.

Pv11 is an insect cell line established from embryos of the African chironomid, *Polypedilum vanderplanki*, an insect known for its ability to undergo anhydrobiosis, a form of extreme desiccation tolerance [8,9]. Similarly, Pv11 cells also display extreme desiccation tolerance: this is induced by treatment with high concentrations of trehalose, a naturally-occurring disaccharide well-known to act in many organisms as a chemical protectant against environmental abiotic stress, including desiccation [10]. Furthermore, trehalose-treated Pv11 cells express the genes associated with desiccation tolerance, such as LEA protein genes [11], which protect biomolecules from desiccation damage [12,13]. We previously showed that induction of desiccation tolerance with trehalose in Pv11 cells allowed effective *in vivo* preservation of exogenously expressed enzymes, where catalytic activity was retained after over one year in the dry state. [14]. These results suggest that the unique stress tolerance of Pv11 cells could be applied to new dry-preservation techniques for biomedical and bioindustrial use.

Exogenous gene expression systems are classified into two main types: constitutive and inducible. Generally, constitutive expression systems employ constitutively active promoters, i.e. the expression of associated genes is not regulated. In Pv11 cells, the 5’-proximal region from −1,269 to +64 bp relative to the transcription start site of the *Pv*.*00443* gene has been identified as one of the strongest constitutive promoters, namely the 121-promoter; in appropriate constructs, this has enabled Pv11 cells to constitutively express several exogenous reporter genes, such as those encoding GFP and luciferase [15]. Constitutive expression systems are useful for the synthesis of recombinant proteins in large quantities; however, excessive overexpression of exogenous proteins can repress cell growth and even cause cell death [16]. In contrast, inducible expression systems produce exogenous proteins only when required. In such systems, even relatively cytotoxic exogenous proteins can be produced, while minimizing their deleterious effects on cell growth. In addition, some inducible expression systems can synthesize recombinant proteins at expression levels comparable to those of major constitutive expression systems [17,18]. These advantages suggest that inducible expression systems should be prioritized for the recombinant production of any protein, regardless of its cytotoxicity. However, inducible expression systems have not yet been developed for Pv11 cells.

One of the most commonly available inducible expression platforms is the Tet-On expression system, which consists of two main factors: a reversible tetracycline-regulated transactivator (rtTA) and a tetracycline-responsive promoter element (TRE) sequence [19]. In the presence of tetracycline or derivatives such as doxycycline (Dox), rtTA is activated, binds to the TRE and subsequently induces the expression of an associated target gene. The TRE is composed of Tet operator (tetO) sequences selectively fused to a minimal promoter suitable for the host cell [20-22].

Minimal promoters are exploited to reduce undesirable leaky expression of recombinant proteins without Dox induction in the Tet-On expression system. In general, a minimal promoter contains DNA regions that recruit the transcriptional initiation machinery, and comprises several functional subregions termed core elements or motifs, such as the TATA box, initiator (Inr), downstream core promoter element (DPE) and bridge [23,24]. These elements and motifs mediate the recruitment of TATA-binding protein (TBP) and TBP-associated factors [25]. Whereas several minimal promoters have been identified and designed for model organisms such as mouse and fruit fly [26], equivalent minimal promoters for most non-model organisms, including *P. vanderplanki*, have yet to be constructed. This represents a bottleneck for the construction of workable Tet-On expression systems for Pv11 cells.

Here we describe the identification of a minimal promoter from a deletion fragment of the 121-promoter of *P. vanderplanki* that is applicable to the Tet-On inducible expression system in Pv11 cells. To establish the proof-of-concept that commercially valuable proteins can be expressed using this approach, we successfully expressed avian myeloblastosis virus reverse transcriptase (AMV RT) using the Tet-On system in Pv11 cells. These results indicate the potential of Pv11 cells for bioindustrial applications.

## 2. Materials and Methods

### 2.1 Cell culture

Pv11 cells were originally established in our laboratory [27]. The cell culture was maintained as described [15]. Briefly, Pv11 cells were cultured using IPL-41 medium (Thermo Fisher Scientific, Waltham, MA) supplemented with 2.6 g/L tryptone phosphate broth (Becton, Dickinson and Company, Franklin Lakes, NJ), 10% (v/v) fetal bovine serum (MP Biomedicals, Santa Ana, CA), and 0.05% (v/v) of an antibiotic and antimycotic mixture (penicillin, amphotericin B, and streptomycin; Merck KGaA, Darmstadt, Germany). As trehalose treatment, the cells were incubated in trehalose mixture (600 mM trehalose (Hayashibara Co., Ltd., Okayama, Japan) containing 10% (v/v) IPL-41 medium).

### 2.2 Vector construction

To construct the pTetO-CMV-AcGFP1 vector, the AcGFP1 gene (Takara Bio, Shiga, Japan) was cloned and amplified using Prime STAR Max Premix (Takara Bio) with specific primers (Suppl. Table 1) from pP121K-AcGFP1 [15] and inserted between the SalI and BglII sites of pTRE3G (Takara Bio) using NEBuilder HiFi DNA Assembly Mater Mix (New England BioLabs, Ipswich, MA). The 202 bp and 266 bp promoters were cloned and amplified with specific primers (Suppl. Table S1) from the pP121K-AcGFP1 series [15] and inserted into the pTetO-CMV-AcGFP1 vector digested with SalI and BglII.

To construct the luciferase Nluc expression vectors, the AcGFP1 genes in the corresponding vectors were replaced by the Nluc gene. The Nluc gene was cloned and amplified using Prime STAR Max Premix (Takara Bio) with specific primers (Suppl. Table 1) from pNL1.1 (Promega, Fitchburg, WI) and inserted between the SalI and BglII sites using NEBuilder HiFi DNA Assembly Master Mix. To construct the promotor-less vector, the 266 bp promoter region of pTetO-266bp-Nluc was replaced with nonsense oligonucleotides generated by annealing the following sequences: 5’-agcttGGCAATCCGGTACTGTTGGTAAAGCCAg-3’as the sense strand; 5’-tcgacTGGCTTTACCAACAGTACCGGATTGCCa-3’ as the anti-sense strand.

pP121K-AcGFP1[15] was modified to make an empty vector by digestion with BamHI and SacII, and replacement of the AcGFP1 sequence with annealed oligonucleotides that introduced new restriction sites for HindIII and XhoI (sense: 5’-gatccaagcttctcgagCCGC-3’, antisense: 5’-GGctcgagaagcttg-3’). We named the empty vector pPv121-MCS (Suppl. Fig. 1 and Suppl. Data). For the pPv121-Tet-On 3G expression vector, pPv121-MCS was digested with BamHI and SacII, and the fragment for Tet-On 3G (the 3^rd^ generation reverse tetracycline-controlled transactivator), which was prepared by PCR with specific primers (Suppl. Table 1), was inserted using NEBuilder HiFi DNA Assembly Master Mix. For the pPv121-Nluc expression vector, pTRE3G (Takara Bio) was digested with XhoI and SalI, and the Nluc fragment, which was prepared by PCR with specific primers (Suppl. Table 1), was inserted using NEBuilder HiFi DNA Assembly Master Mix. For construction of the luc2-reference vector, a 632 bp fragment of the 121-promoter [15] was amplified by PCR (Suppl. Table 1) and inserted between the BglII and HindIII sites of pGL4.10 (Promega) using NEBuilder HiFi DNA Assembly Master Mix.

**Figure 1.**
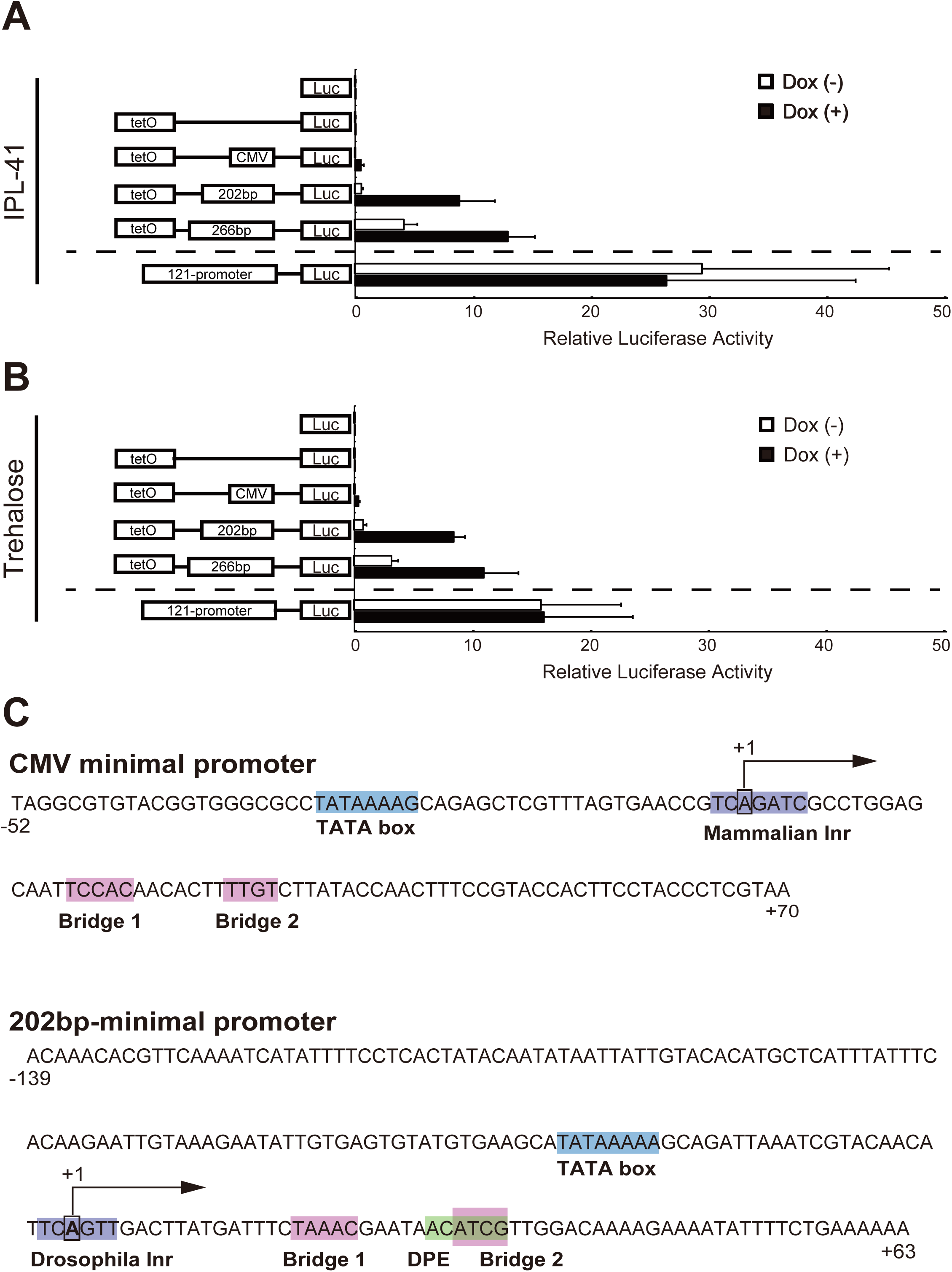
Identification of a minimal promoter for an inducible expression system in Pv11 cells. Luciferase activity was measured in vector-transfected Pv11 cells cultured under standard conditions (IPL-41 medium; A) or with 600 mM trehalose treatment (B). A scheme of vector constructs is shown on the left and relative values in the luciferase assay are shown on the right. Normalized values are expressed as means ± standard deviations (SD). (C) The predicted core promoter elements of the CMV and 202 bp minimal promoters. The core promoter prediction program, the Elements Navigation Tool, was used with a default setting.

A DNA fragment for the AMV RTα gene was synthesized by Eurofins Genomics K.K, Tokyo, Japan. The AMV RTα gene was then amplified by PCR with specific primers (Suppl. Table 1) and inserted into pPv121-MCS digested with BamHI and SacII using NEBuilder HiFi DNA Assembly Mater Mix. For the pTetO-202bp-AMV RTα expression vector, the AMV RTα sequence was amplified and inserted into pTetO-202bp-Nluc digested with SalI and BglII using NEBuilder HiFi DNA Assembly Master Mix.

### 2.3. Transfection and luciferase reporter assay

The cells used in each experiment were seeded at a density of 3 × 10^5^ cells per mL into a 25 cm^2^ cell culture flask and grown at 25°C for 6 days before transfection. Transfection into Pv11 cells was carried out using a NEPA21 Super Electroporator (Nepa Gene, Ichikawa, Chiba, Japan) as described previously [28]. For transient expression of luciferase, a mixture of 2 μg of either Nluc-reporter vector or 121-Nluc expression vector, 10 μg luc2-reference vector and 0.5 μg Tet-On 3G expression vector was transfected into the cells. One day after transfection, the medium was replaced with IPL-41 or trehalose mixture with or without doxycycline (Dox) at 1 μg/mL. The cells were collected 24 h later, and luciferase activity was measured using an ARVO luminometer (PerkinElmer, Waltham, MA) with the Nano-Glo Dual-Luciferase Reporter Assay System (Promega).

### 2.4. Prediction of core promoter elements

To identify putatively functional motifs in the 202 bp minimal promoter, the Elements Navigation Tool (http://lifefaculty.biu.ac.il/gershon-tamar/index.php/element-description) [29] was used with default settings.

### 2.5. GFP expression using the Tet-On system

For transient expression of GFP, three combinations of the corresponding vectors were transfected into Pv11 cells as follows: 10 μg pPv121-MCS; 1 μg pPv121-Tet-On 3G and 9 μg pTetO-202bp-AcGFP1; 1 μg pPv121-MCS and 9 μg pPv121-AcGFP1. One day after transfection, the medium was replaced with IPL-41 or trehalose mixture with or without Dox at 1 μg/mL. After 24 h, the AcGFP1 fluorescence in the cells was viewed using a Biozero BZ-X700 microscope (Keyence, Osaka, Japan). AcGFP1 expression was evaluated by Western blot analysis.

### 2.6. AMV RTα expression using the Tet-On system

For transient expression of AMV RTα, three combinations of the vectors were transfected into Pv11 cells as follows: 10 μg pPv121-MCS; 1 μg pPv121-Tet-On 3G and 9 μg pTetO-202bp-AMV RTα; 1 μg pPv121-MCS and 9 μg pPv121-AMV RTα. One day after transfection, the same treatment was performed as described above. Twenty-four hours after the treatment, AMV RTα expression in the cells was evaluated by Western blot analysis, and reverse transcriptase (RT) activity was measured.

### 2.7. Antibodies and Western blot analysis

As primary antibodies, monoclonal anti-GFP antibody (M048-3, Medical & Biological Laboratories, Nagoya, Japan; dilution ratio: 1/1,000) and monoclonal anti-His-tag antibody (D291-3, Medical & Biological Laboratories; dilution ratio:1/2,000) were used to detect AcGFP1 and AMV RTα, respectively. Cells were lysed with RIPA lysis buffer (Nacalai Tesque, Kyoto, Japan) for 30 min at 4°C. After centrifugation, aliquots of the supernatant were subjected to protein quantification with a Pierce BCA Protein Assay Kit (Thermo Fisher Scientific) and SDS-PAGE using a TGX Stain-Free FastCast Acrylamide Kit (Bio-Rad, Hercules, CA). For AcGFP1 detection, 3 μg protein/lane was used, while for AMV RTα detection, 15 μg protein/lane was used for SDS-PAGE. After transferring to a PVDF membrane using a Trans-Blot Turbo (Bio-Rad), the membrane was blocked with 1% skimmed milk in TBS with 0.1% Tween 20 (TBST) at room temperature for 1 h. Primary antibodies were diluted in Can Get Signal Solution1 (TOYOBO, Osaka, Japan), and incubated at room temperature for 1 h. The membrane was washed three times with TBST for 10 min each wash, followed by incubation of HRP-conjugated secondary antibodies (goat anti-mouse IgG, 81-6720, Thermo Fisher Scientific; dilution ratio: 1/2000) in Can Get Signal Solution 2 (TOYOBO) at room temperature for 1 h. The membrane was washed three times with TBST (10 min each wash), and chemiluminescent signals from ECL Prime detection reagents (Cytiva, Little Chalfont, Buckinghamshire, UK) were captured on a ChemiDoc™ Touch imaging system (Bio-Rad).

### 2.8. Reverse transcriptase assay

Cells were lysed with a PBS (Merck KGaA)-based lysis buffer containing 0.5% v/v Nonidet P-40, 10% w/v glycerol, 5mM DTT and Complete Mini EDTA-free Protease Inhibitor Cocktail (Merck KGaA) and sonicated for 30 s (UR-21P, TOMY SEIKO, Tokyo, Japan). After centrifugation, aliquots of the supernatant were diluted with nuclease-free water and subjected to DNA determination using a Qubit 2.0 fluorometer and a Qubit dsDNA HS quantitation assay (Thermo Fisher Scientific). All samples were diluted to 0.436 ng/μL with lysis buffer and subjected to a reverse transcriptase assay (Thermo Fisher Scientific). Recombinant AMV RTα (Nippon Gene, Tokyo, Japan) was used to prepare a standard curve. The samples and recombinant AMV RTα were diluted with lysis buffer, then mixed with a reaction mixture and incubated at 25°C. After 1 h incubation, the RT reactions were stopped by adding EDTA, and GFP fluorescence from PicoGreen was quantified using an ARVO luminometer (PerkinElmer).

### 2.9. Statistical analysis

All data were expressed as mean ± SD. Statistical significance among more than two groups was examined by ANOVA followed by a Tukey post-hoc test (Figs. 1A, 3B, Suppl. Tables 2 and 3). A *p*-value < 0.05 denoted a statistically significant difference. GraphPad Prism 8 software (GraphPad, San Diego, CA) was used for the statistical analyses.

## 3. Results

### 3.1. Identification of a minimal promoter for the development of an effective inducible expression system in Pv11 cells based on the Tet-On system

To develop an effective inducible expression system, we first attempted to identify an efficient minimal promoter in Pv11 cells. We previously suggested that deletion fragments of the 121-promoter, including fragments 266 bp and 202 bp in length, might be candidates for a minimal promoter because such fragments have low activity in a luciferase assay [15]. Thus, we carried out a luciferase assay using vectors harboring either one of these candidates, or the cytomegalovirus (CMV) minimal promoter, or no promoter, fused with the tetO sequence to evaluate whether these fragments could work as a minimal promoter in Pv11 cells. The pPv121-Nluc vector containing the fully functional 121-promoter was used as a positive control for this assay. As reported for other insect cells [30,31], the CMV minimal promoter did not promote luciferase activity in the presence of Dox in Pv11 cells (Fig. 1A and Suppl. Table 2). In contrast, under standard culture conditions without Dox treatment, the 266 bp fragment resulted in leaky expression of luciferase, while the 202 bp fragment showed deficient transcriptional activity comparable to that of negative controls, such as pNluc, pTetO-Nluc and pTetO-CMV-Nluc (Fig. 1A and Suppl. Table 2). Dox treatment enhanced luciferase expression in cells transfected with the vectors harboring the 266 bp and 202 bp fragments (3- and 16-fold induction, respectively) (Fig. 1A and Suppl. Table 2). These results indicate that, of the two fragments tested, the 202 bp-fragment functions best as a minimal promoter in the Tet-On inducible system in Pv11 cells.

Pv11 cells display extreme desiccation tolerance when induced by highly concentrated (600 mM) trehalose solution [8]. Trehalose treatment is known to trigger specific transcriptional regulatory networks that activate anhydrobiosis-related genes [32], suggesting that trehalose might affect the expression level of the exogenous Tet-On inducible system. Thus, we conducted luciferase assays for the corresponding vectors in trehalose-treated Pv11 cells. The results showed that trehalose treatment did not interfere with the inducibility of the Tet-On system in Pv11 cells (Fig. 1B and Suppl. Table 2). In particular, Dox treatment of Pv11 cells transfected with the vector harboring the 202 bp fragment augmented luciferase expression 11-fold (Fig. 1B). Therefore, in Pv11 cells, the Tet-On inducible system including the 202 bp-fragment (Fig. 1C) as a minimal promoter can be used both during normal culture and during induction of desiccation tolerance by trehalose.

The 202 bp minimal promoter fragment contained functional core elements, such as TATA box, Inr, DPE and bridge elements, similarly to the CMV minimal promoter (Fig. 1C), although the nucleotide sequences of these elements were different in each promoter. This suggests that mammalian promoters, such as the CMV minimal promoter, may not function in Pv11 cells due to sequence differences in core promoter elements.

### 3.2. AcGFP1 expression using the Tet-On system in Pv11 cells

To assess the versatility of the Tet-On system, we expressed the AcGFP1 gene in Pv11 cells. We tested this with the pTetO-202bp-AcGFP1 vector, constructed as described in the Methods, using pPv121-MCS (empty vector) and pPv121-AcGFP1 as negative and positive controls, respectively. AcGFP1 fluorescence was observed in Dox-treated cells transfected with pTetO-202bp-AcGFP1 vectors under both normal culture (IPL-41 medium) and trehalose treatment conditions. In contrast, only a few AcGFP1-positive cells were observed in the absence of Dox (Fig. 2A). Western blotting showed a massive accumulation of AcGFP1 protein in Dox-treated groups transfected with pTetO-202bp-AcGFP1 vector under both IPL-41 medium and trehalose treatment conditions, while almost no AcGFP1 protein was detected in the absence of Dox (Fig. 2B). Together with the luciferase assay data (Fig. 1A and B), these results show that the Tet-On system with the 202 bp minimal promoter can be used for the inducible expression of reporter genes, including GFP and luciferase, in Pv11 cells.

**Figure 2.**
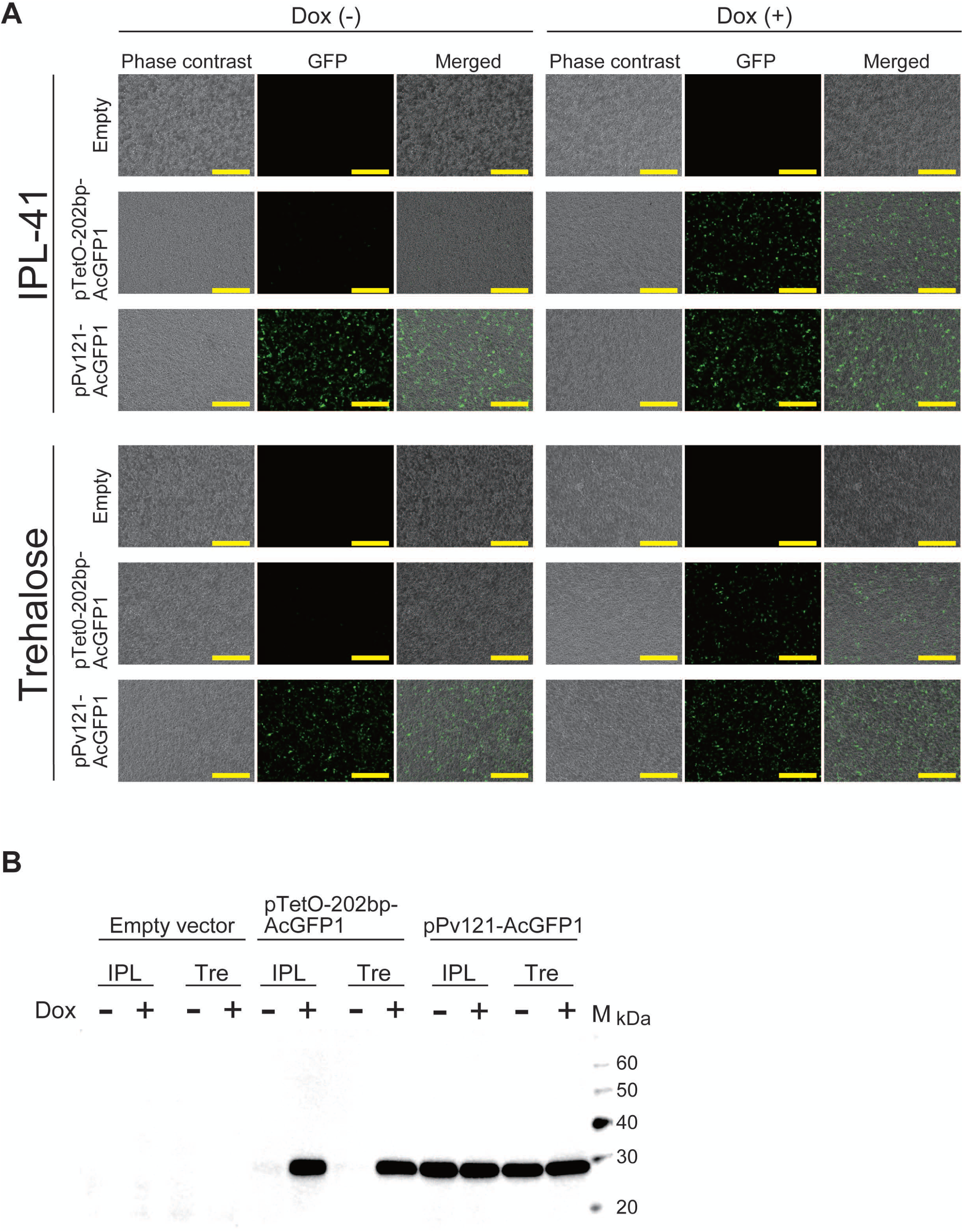
AcGFP1 expression using the Tet-On system in Pv11 cells. (A) The images of the cells were acquired using a BZ-X700 fluorescence microscope. The upper and lower images show the cells cultured in IPL-41 medium and 600 mM trehalose solution, respectively. Scale bars, 200 μm. (B) Western blot analysis was performed on extracts of Pv11 cells transfected with the indicated vectors. IPL, IPL-41 medium; Tre, 600 mM trehalose solution; Lane M, protein marker.

### 3.3. AMV RTα expression and measurement of RT activity

To examine whether Pv11 cells, like other insect cells, can express functional proteins with complex structures, such as proteins containing multiple domains, we focused on AMV RT, which is one of the most widely used RNA-dependent DNA polymerases for medical diagnoses [33] and basic research. AMV RT is a heterodimer consisting of a 63 kDa α-subunit and a 95 kDa β subunit [34], but the α-subunit alone has RT activity [35]. Therefore, we chose AMV RTα to establish the proof-of-concept that commercially valuable proteins can also be expressed in Pv11 cells.

Accordingly, we prepared the pTetO-202bp-AMV RTα vector as described in the Methods, and transfected this into Pv11 cells, alongside pPv121-MCS and pPv121-AMV RTα as negative and positive controls, respectively. As shown in Figure 3A, AMV RTα was observed to accumulate under Dox control for the Tet-On vector both in IPL and trehalose culture conditions, while no AMV RTα protein was detected with the empty vector. On the other hand, transfection with the pPv121-AMV RTα vector allowed the production of AMV RTα protein independently of the medium or the presence of Dox (Fig. 3A). Next, RT activity was assessed. As shown in Figure 3B, the RT activity was significantly higher in Dox-treated cells than in untreated counterparts using the Tet-On expression system, while the constitutive expression vector yielded high RT activity independently of culture conditions (Fig. 3B and Suppl. Table 3). These results clearly show that Pv11 cells can produce a large and complex enzyme, such as AMV RTα, using the Tet-On inducible expression system.

**Figure 3.**
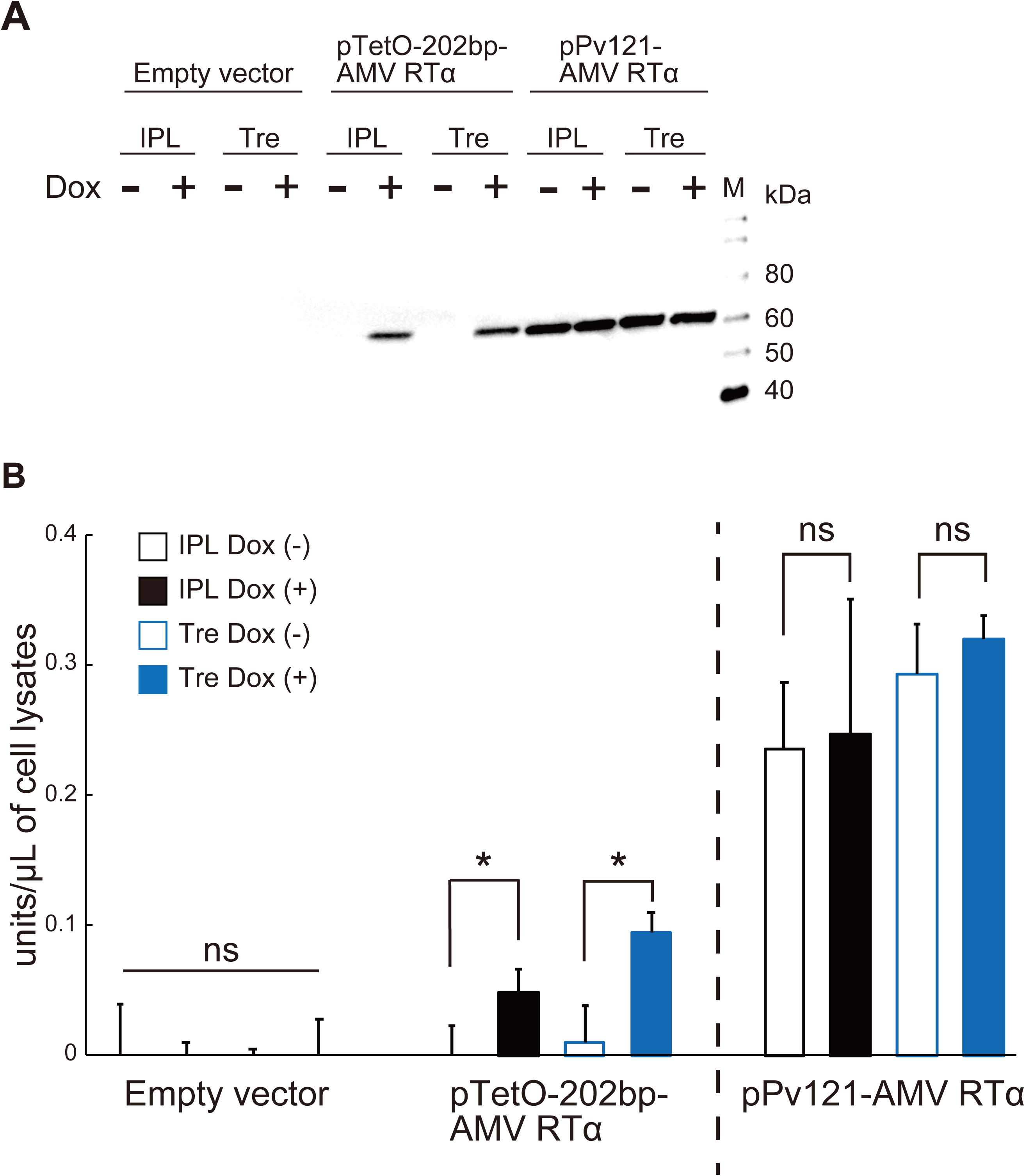
AMV RTα expression and direct measurement of reverse transcriptase activity. (A) Western blot analysis was performed on extracts of Pv11 cells transfected with the indicated vectors. IPL, IPL-41 medium; Tre, 600 mM trehalose solution; Lane M, protein marker. (B) AMV RTα activities were measured directly in lysates of Pv11 cells transfected with the indicated vectors. Normalized values are expressed as mean ± standard deviation (SD). *, significant at *p<*0.05; ns, not significant; n=3 in each group.

## 4. Discussion

Although a constitutive expression system was previously established in the anhydrobiotic cell line, Pv11 [15], no inducible system has been available so far. The present study identified a minimal promoter for Pv11 cells and applied it in the Tet-On inducible expression system. Luciferase and GFP proteins were successfully and inducibly expressed in this novel system under both regular and trehalose-treatment culture conditions. Furthermore, we showed that AMV RTα protein could be inducibly expressed in Pv11 cells without losing its RT activity. This highlights the potential of Pv11 cells for the expression of larger and more complex proteins than reporter proteins and shows that Pv11 cells could have novel biotechnological applications.

The establishment of a Tet-On inducible expression system in Pv11 cells could accelerate the clarification of the molecular mechanisms underlying anhydrobiosis in this cell type and more generally. For example, the Tet-On expression system has been used for inducible gene knockdown or knockout [36-40], thereby providing valuable insights into specific gene functions. Such tightly regulated loss-of-function experiments will be helpful in examining whether or not a gene contributes to anhydrobiosis in Pv11 cells. Furthermore, minimal promoters constitute an essential component of massively parallel reporter assays [41-44], which are among the most powerful genome-wide screening systems. Thus, combining these cutting-edge molecular biological technologies with our current findings should result in a markedly improved understanding of the mechanisms of anhydrobiosis in Pv11 cells.

Several studies have shown that the mammalian CMV and SV40 promoters do not function in insect cell lines derived from *Drosophila* and *Spodoptera* species [30,31]. In the present study, we found that the CMV minimal promoter also does not function in Pv11 cells (Fig. 1A and B). This result can probably be explained by sequence differences in the core promoter elements. The Inr in the *P. vanderplanki* 202 bp-promoter is TCA_+1_G**<u>T</u>**T, whereas in the CMV minimal promoter the Inr is TCA_+1_G**<u>A</u>**TC (Fig. 1C). Indeed, the consensus sequence for the Inr is YYA_+1_N**<u>W</u>**YY for humans while it is YYA_+1_K**<u>T</u>**Y for *Drosophila* [23]. Clearly, the selection of a promoter with sequence elements that function in the target host cell is essential; in this context, the use of an endogenous promoter is particularly appropriate.

To develop the inducible expression system described here, we needed to define a minimal promoter and demonstrate its application. Previously, we identified 266 bp, 202 bp, and 137 bp fragments from the *P. vanderplanki* 121-promoter as candidate minimal promoters for Pv11 cells [15]. We tested the first two of these fragments, and found the 202 bp fragment to perform well in the Tet-On system. However, it was difficult to obtain sufficient quantities of the plasmid harboring the 137 bp fragment for expression experiments (Suppl. Figs. 2A and B), probably due to replication interference caused by its stem-loop structure (Suppl. Fig. 2C). Thus, the 137 bp fragment was excluded from our experiments.

Several studies have shown that the maximum transcriptional activity of the Tet-On inducible expression system is comparable to that of constitutive counterparts [18]. In this study, however, the induced Nluc activity of the Tet-On system was only about 30-50% of that of the 121-promoter under IPL (8.88±3.05 relative luciferase activity (RLA) vs. 26.50±16.05 RLA) and trehalose (8.45±0.99 RLA vs. 16.14±7.57 RLA) culture conditions (Figs. 1A and B). This is also reflected in the different relative AMV RTα activities observed between the constitutive and Tet-On inducible expression systems (Fig. 3B). These results probably reflect the need to co-transfect the Tet-On 3G expression and tetO reporter vectors, both of whose expression are controlled by tetO sequences, while only one vector is sufficient to express the gene-of-interest in the constitutive expression system. In particular, in our experiments we used two vectors (e.g. pPv121-Tet-On 3G and pTetO-202bp-Nluc) for the Tet-On expression system and one vector (pPv121-Nluc) for the constitutive expression system. To reduce the complexity of the Tet-On system, and thus potentially improve expression levels, we are attempting to establish Pv11 cells that stably express the Tet-On 3G transactivator protein.

For industrial protein production, the bacterium *Escherichia coli* is one of the most extensively used prokaryotic cells; however, eukaryotic cells must be used instead when the protein-of-interest involves complex post-translational modifications and/or has multiple structural domains that are essential for its function [45,46]. Indeed, the expression of eukaryotic multi-domain proteins in *E. coli* often results in non-functional products due to differences in folding machinery and post-translational modifications between prokaryotes and eukaryotes [45,46]. Additionally, it is known that certain RT variants cannot be expressed as active or soluble heterodimers in *E. coli* without additional expression of appropriate molecular chaperones [47,48]. Indeed, in our experience (unpublished results), AMV RT, which is a multi-domain protein (comprising finger, palm, thumb, connection and RNase H domains) [35], is difficult to obtain in a soluble form in sufficient quantities in *E. coli*. In contrast, we demonstrated in the present study that AMV RTα can be expressed in Pv11 cells without losing its specific activity. This result suggests that it should be possible to express a variety of proteins, including large multi-domain proteins, in Pv11 cells.

We used a simplified method here for detecting AMV RTα activity from crude cell lysates. Although a certain amount of nonspecific background fluorescence from the lysates of Pv11 cells cannot be avoided, this method enabled us to decrease sample volumes markedly: transfected cells were harvested from individual wells of a 48-well plate, and only 1 µL of each lysate was required for the enzyme assay. Although further improvements in our method may be necessary, it could be adapted for easy screening of useful AMV RT mutants. We will also examine the possibility of preserving exogenously expressed AMV RTα in dried Pv11 cells.

In conclusion, the identification of a minimal promoter for Pv11 cells led to the establishment of a Tet-On inducible expression system in Pv11 cells. We showed that, in addition to the reporter proteins luciferase and AcGFP, AMV RTα could be expressed successfully in the Pv11 Tet-On system without losing its RT activity. These results show that Pv11 cells have significant potential for bioindustrial protein production.

## Acknowledgements

We are grateful to Tomoe Shiratori for routine maintenance of Pv11 cells.

## Figures legends

**Supplementary Figure 1.**
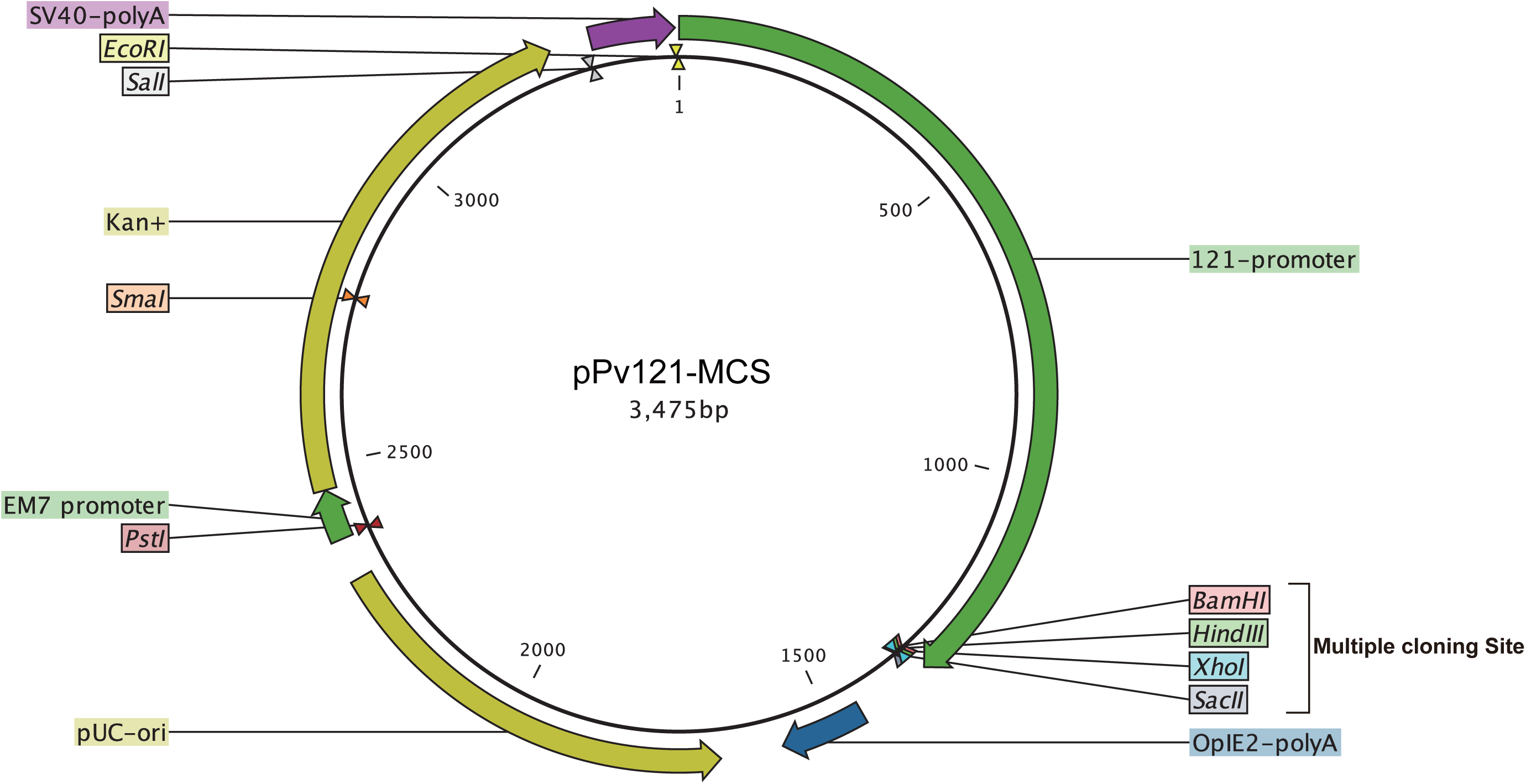
Map of pPv121-MCS vector. The multiple cloning site of the vector comprises BamHI, HindIII, XhoI and SacII sites.

**Supplementary Figure 2.**
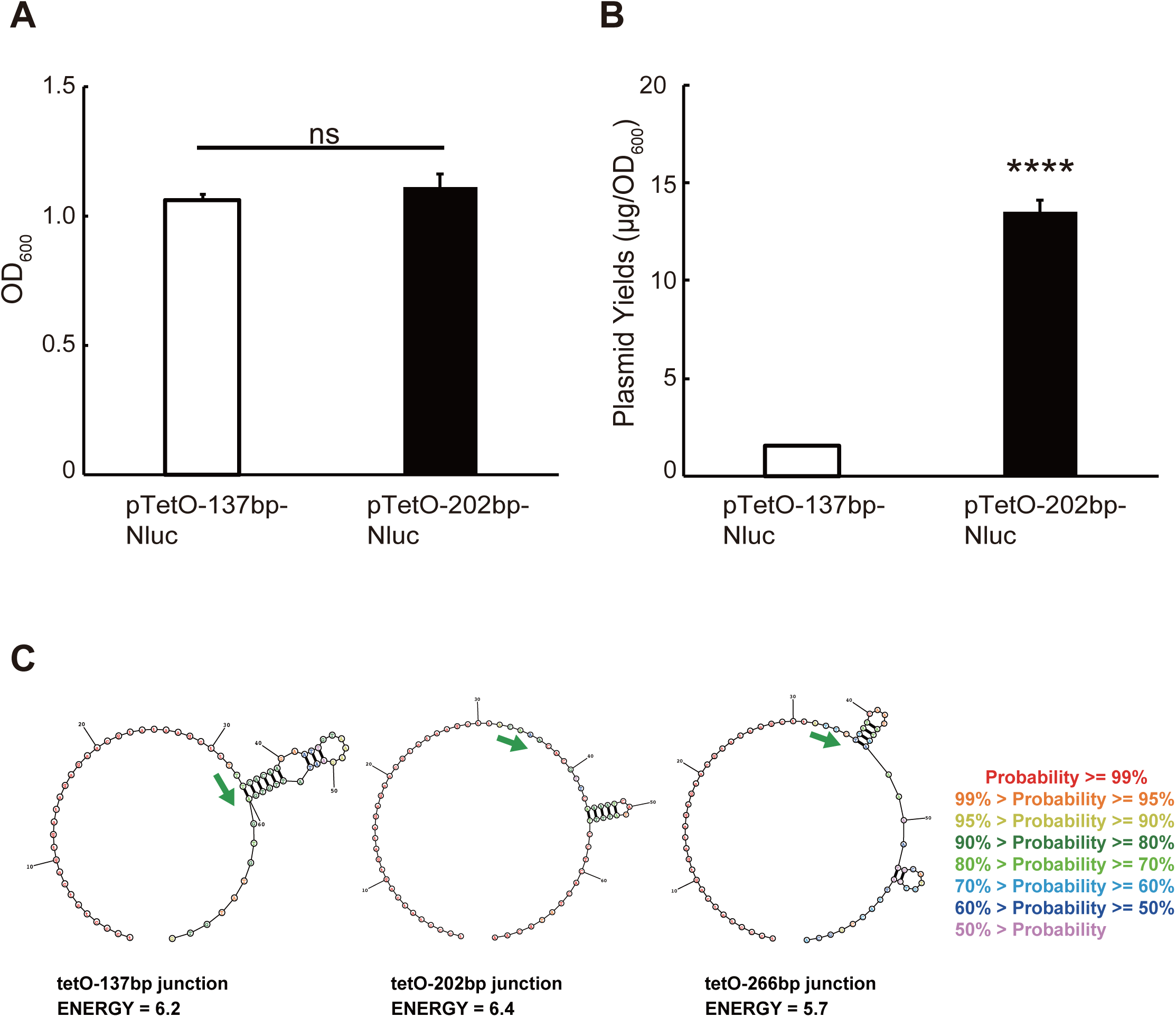
The 137 bp-fragment from the 121-promoter and its secondary structure. (A) Optical density (OD) of *E. coli* transformed with pTetO-137bp-Nluc and pTetO-202bp-Nluc. The transformants were cultured in LB medium containing ampicillin at 37°C for 16 h, then OD_600_ was measured. (B) Yields of the pTetO-137bp-Nluc and pTetO-202bp-Nluc plasmids. The plasmids were purified, and the concentration was measured. Normalized values are expressed as means ± standard deviations (SD). \*\*\*\**p*<0.0001; ns, not significant; n=3 in each group. (C) Secondary structure of 69 bp junction regions of the tetO area (31 bp) and 137 bp, 202 bp and 266 bp fragments. The secondary structures of tetO and each fragment region were predicted using the MaxExpect algorithm (Reuter & Mathews, 2010) for DNA sequences with the remaining parameters set to default. Green arrows define the starting point for each fragment region. The tetO area has no possible secondary structures, while the sequences of the various fragments tested may fold into different stem-loop structures, which might affect plasmid function melting to single strands, as in replication. According to free energy values, the tetO 137 bp stem-loop is the most stable. Reference: Reuter, J.S. and Mathews, D.H. “RNA structure: software for RNA secondary structure prediction and analysis.” BMC Bioinformatics, 11:129. (2010)

**Supplementary Table 1.**
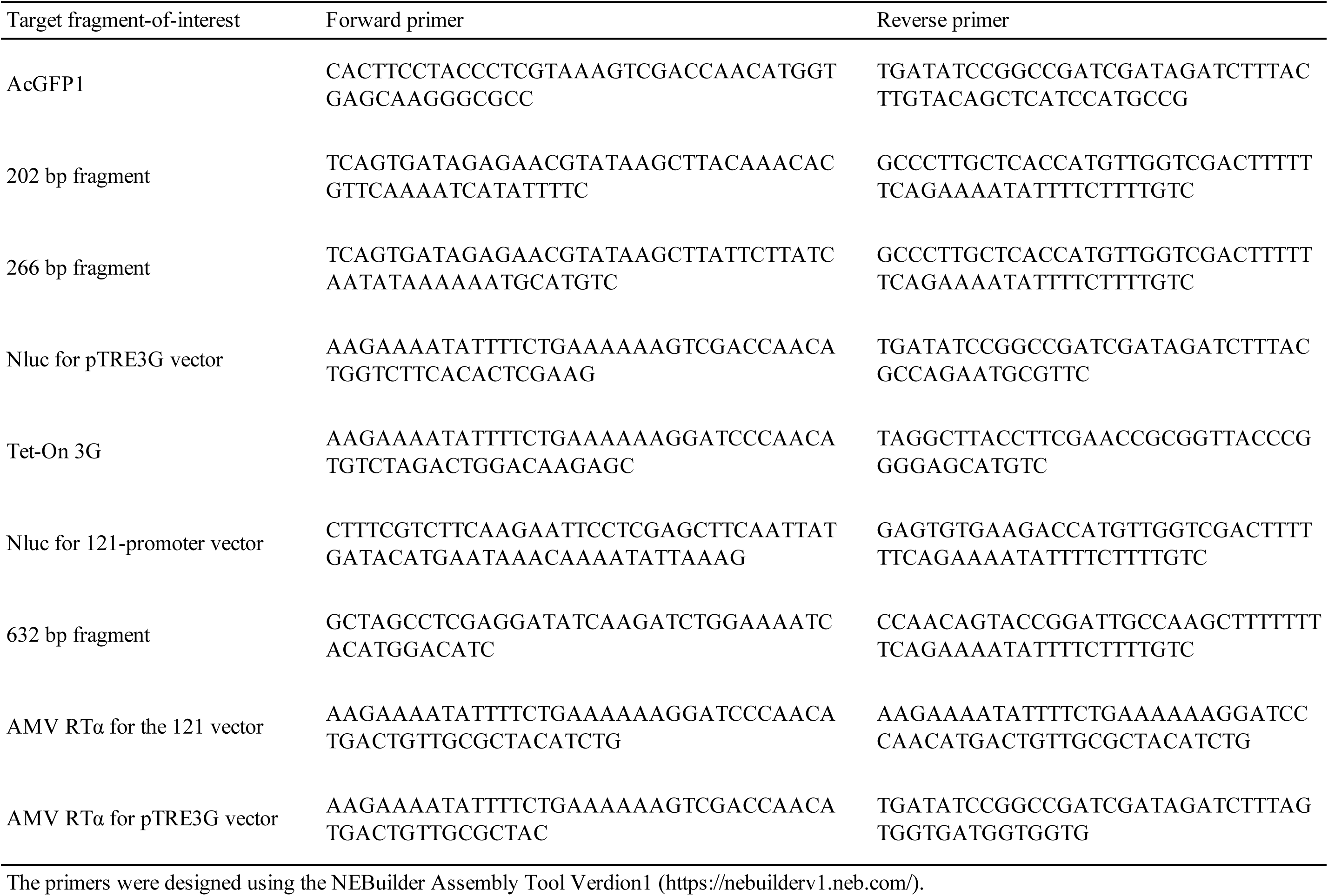
Primers for the construction of AcGFP1, Nluc and AMV RTα expression vectors.

**Supplementary Table 2.**
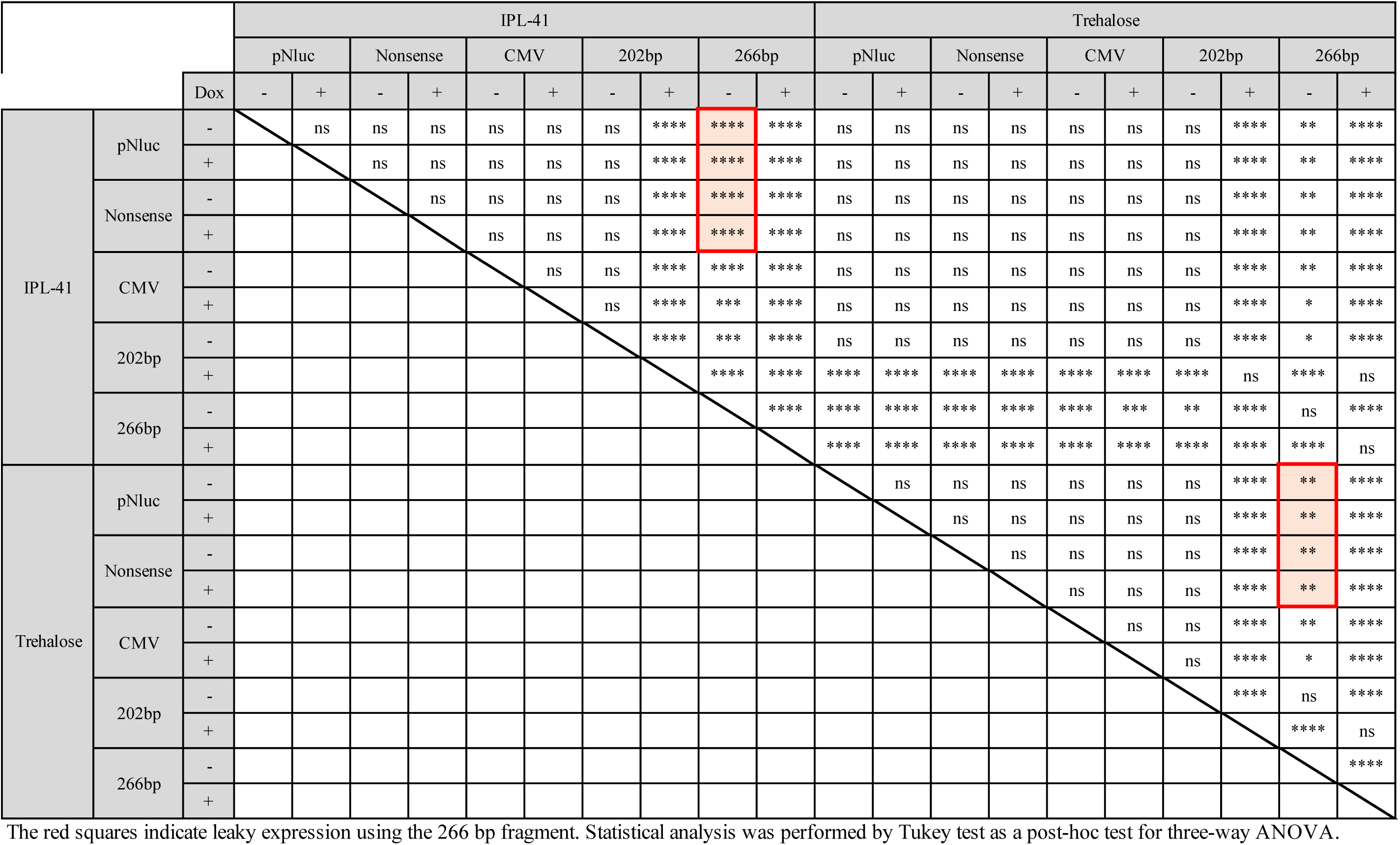
Statistical analysis of luciferase activities of cells transfected with the corresponding vectors in Fig. 1A.

**Supplementary Table 3.**
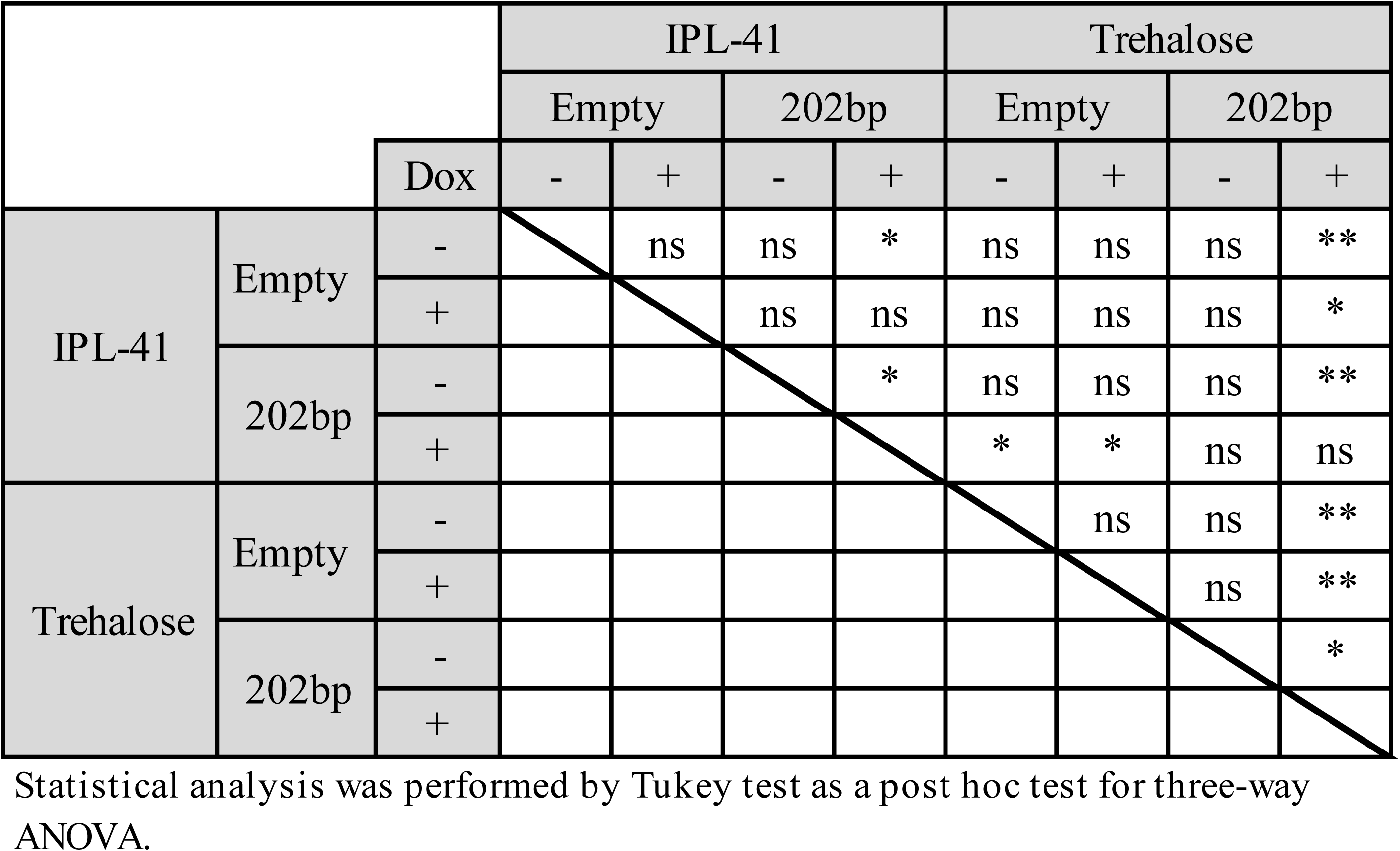
Statistical analysis of reverse transcriptase activities of cells transfected with the corresponding vectors in Fig. 3B.

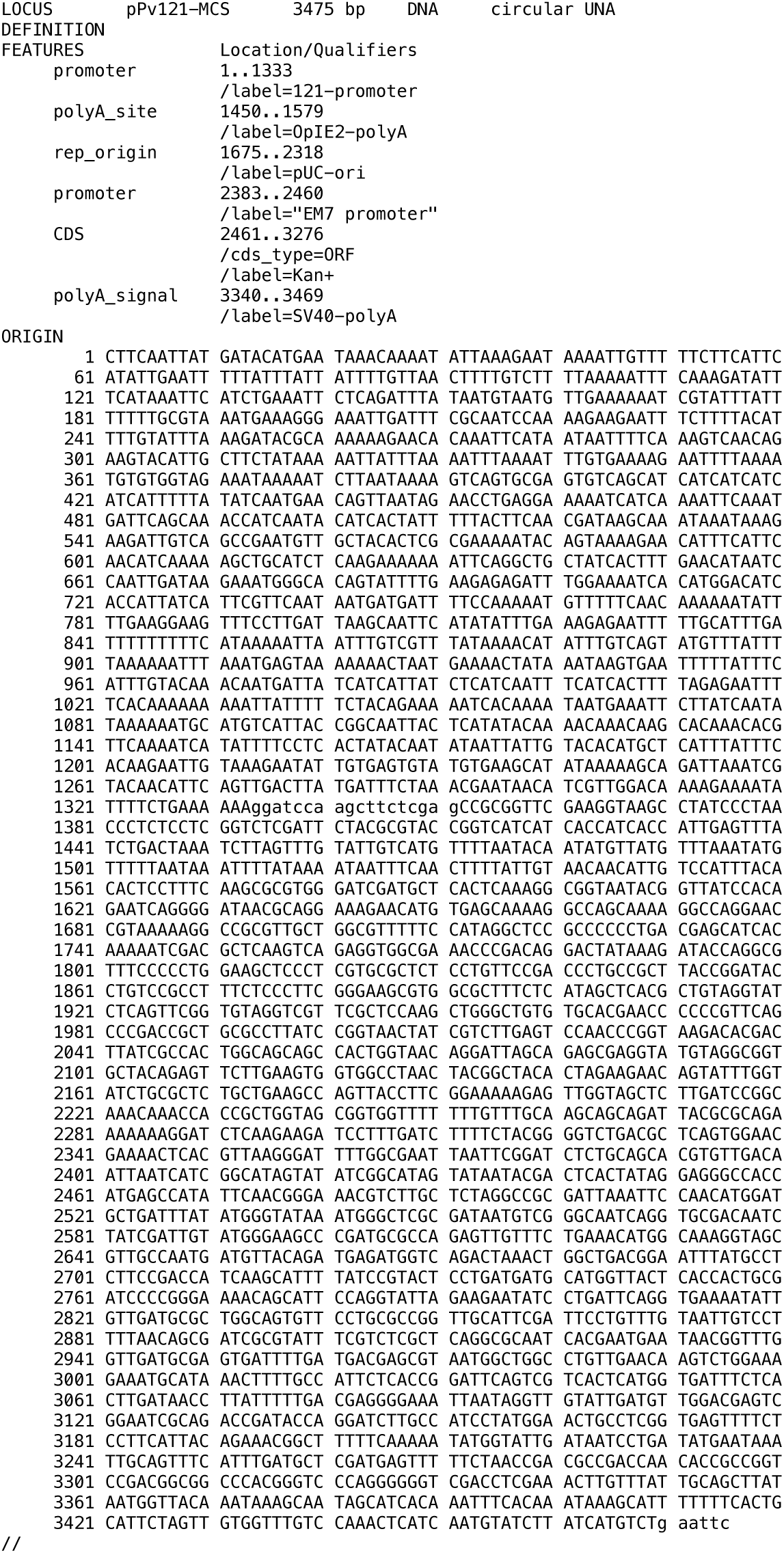

